# The Interplay of Resource Availability and Parent Foraging Strategies on Juvenile Sparrow Individual Specialization

**DOI:** 10.1101/2024.02.29.582146

**Authors:** Natalia Ricote, Constanza Weinberger, Natalia Ramírez-Otarola, Sara Bustamante, María Lucía Málaga, Gonzalo Barceló, Pablo Sabat, Seth D. Newsome, Karin Maldonado

## Abstract

Temporal variation in resource availability, amplified by global change, may have strong impacts on species breeding at temperate and high latitudes that cue their reproduction to exploit seasonal resource pulses. This study examines how resource availability and parental care influence niche partitioning between and within age classes in the rufous-collared sparrow, which provides extensive parental care. We hypothesized juveniles would exhibit narrower niches focused on high-quality resources compared to adults, regardless of resource availability. We used stable isotope analysis to quantify individual and population niches in juveniles and adults across the breeding season in two cohorts experiencing contrasting resource landscapes. Contrary to our initial hypothesis, juveniles exhibited greater among individual diet variation and smaller total niche widths (i.e., higher levels of individual specialization) during periods of high food availability in comparison to periods of food scarcity. Interestingly, total niche width and individual specialization of adults remained stable across seasons despite a shift in trophic level, highlighting their potential role in providing a consistent diet for their young. These findings reveal a dynamic interplay between resource availability, parental care, and individual specialization, with important implications for understanding population resilience under variable resource scenarios. The study also suggests that adult sparrows modify their provisioning strategies based on resources, potentially buffering offspring from environmental fluctuations. Understanding age-specific responses to resource variation is crucial for predicting species responses to ecological conditions, particularly in regions like central Chile where seasonal resource limitation is expected to become more variable in response to climate change.

## Introduction

Global change is expected to increase fluctuations in the availability of resources (Jia et al., 2019), and within-population variability in resource use may increase its resilience to such environmental changes (Ramos et al., 2020; Riverón et al., 2021). Diet variation within a population can arise from individuals belonging to different age classes, sex, or discrete morphs (Bolnick et al. 2003). Additionally, continuous morphological, physiological, and behavioral differences among individuals, independent of these categories, can also influence dietary choices (Araujo et al. 2011). This results in individuals using a narrower range of resources compared to the overall population (Costa-Pereira et al., 2018). In other words, while the population as a whole may exploit a broad spectrum of resources, individual members may focus on a more limited subset (Maldonado et al. 2019). This variation in resource use within a population, known as individual specialization (IS), can be driven by mechanisms similar to those that cause resource partitioning among different species, which serve to reduce niche overlap and intraspecific competition (Keast, 1977; Maiorana, 1978; Svanbäck and Bolnick, 2005). Furthermore, IS can lower the costs of switching between food items and allow for the efficient exploitation of a narrow range of resources, while generalists rely on diverse resources and often exploit each one with less efficiency (Araújo et al., 2011; Bolnick et al., 2011; Maldonado et al. 2019).

In species with parental care, individuals need to compete for resources not only for themselves but also to provision their offspring (Beaulieu and Sockman, 2014; Arco et al., 2022). The initial assumption is that parents feed their offspring a diet as diverse as what they consume, leading to similar resource use and no niche partitioning between adults and juveniles. However, in birds, parental decisions about how to allocate food to their nestlings are strongly influenced by nestling behavior, parental sex, life history traits, and the nutritional needs of their offspring, which require high-quality food to rapidly grow and enhance their probability of survival (Alonso et al., 2012; Raya Rey et al., 2012; Beaulieu and Sockman, 2014). Indeed, it has been suggested that protein intake is higher at early ages due to the predominance of anabolic reactions caused by the formation of new tissues (Kyriazakis and Oldham, 1993). This is consistent with studies on the overlap of population niches among different age classes, which show that juveniles occupy higher trophic levels compared to adults (Hobson, 1993; Hammerschlag-Peyer et al., 2011). Balancing self-maintenance with nurturing offspring becomes particularly challenging when resources are scarce (Arco et al., 2022). Therefore, parents’ foraging decisions, which shape niche characteristics, are crucial for the reproductive success and survival of both themselves and their offspring (Stearns, 1998; Arco et al., 2022). Variation in resource availability has a well-documented influence on niche breadth, niche partitioning, levels of IS, and may also affect trophic level in omnivorous species (Finke and Snyder, 2008; Araújo et al., 2011; Stephens et al., 2019). Several studies have reported niche partitioning between adults and juveniles, as well as differences in IS among age classes due to disparities in foraging abilities, morphology (*e.g.,*body mass, bill length), or social status (Sword and Dopman, 1999; Durell, 2003; Frédérich et al., 2010; Vander Zanden et al., 2013; Hall et al., 2021). Nevertheless, a common feature in these studies is that they report differences between adults and juveniles in species that have little or no parental care or in juveniles that can forage independently (Durell, 2003; Hall et al., 2021). Moreover, these studies have not considered resource variation as a potential explanatory variable in niche differentiation within population (Hall et al., 2021).

In altricial, central forager, short-lived bird species with parental care -such as small passerines-, we expect parents to select high-quality food to ensure their brood’s survival even if it results in more competition with other adults during periods of low food availability (Beaulieu and Sockman, 2014). Thus, juveniles should exhibit narrower niches focused on high-quality prey occupying higher trophic levels with high protein contents. As a result, this would lead to lower degrees of individual specialization compared to adults, regardless of environmental shifts in resource availability. On the contrary, adults are expected to have higher levels of IS during conditions of high resource availability (Hall et al., 2021). No studies have examined how parental care, specifically the provisioning of food from parents to offspring, affects niche partitioning both between and within age classes under contrasting levels of resource availability.

To assess these hypotheses, we studied the rufous-collared sparrow (*Zonotrichia capensis*) that provides parental care while chicks are in the nest and during the post-fledging period as juveniles learn to forage independently (Miller and Miller, 1968; Tubaro, 1990). To compare niche partitioning between and within age classes, we utilized the species’ ability to generate multiple broods within a single breeding season, which allowed two distinct cohorts of nestlings and adults to experience contrasting resource availability. We used isotope-based estimates of dietary variation, an effective approach to quantify individual specialization by analyzing individual and population isotopic variance in consumer tissues that reflect diet over different timescales (Matich et al., 2011; Hanson et al., 2015; Bond et al., 2016; Maldonado et al., 2017). Nitrogen isotope (δ^15^N) values have been widely used to characterize trophic levels and food-chain length (Post, 2002). In this study, we measured δ^15^N values in blood plasma, red blood cells, and claw tips collected from each individual as proxies for dietary variation to estimate individual specialization and trophic levels of juveniles and adults throughout the breeding season (Bearhop et al., 2004; Vander Zanden et al., 2010).

## Materials and Methods

### Fieldwork and Tissue Collection

The study site, Quebrada de la Plata (33°31’S 70°50’W) in central Chile, is characterized by a Mediterranean climate with cold wet winters and relatively hot dry summers (Di Castri and Hajek, 1976). The average accumulated winter precipitation is 230 mm, and the during the breeding season in our study, it was 180 mm (CR2, 2024). In addition, at this site the availability of insects and plants varies seasonally being highest in spring and lowest in winter; summer values also are lower than in spring (Lopez-Calleja, 1995). We captured rufous-collared sparrows with mist nets in spring (November 2020) and summer (January 2021). We categorized captured individuals into adults (N_spring_= 16, N_summer_= 20) or juveniles (fledglings; N_spring_= 12, N_summer_= 8) according to plumage characteristics described in Miller and Miller (1968). We ringed the captured birds to ensure the independence of our samples, avoiding the resampling of the same individuals across seasons.

Parental care in this species extends from hatching until complete independence at 40 days old, with both parents actively feeding the chicks. Fledglings leave the nest after 18-20 days; at which time they start learning to eat on their own and can feed independently from four weeks old. Thus, they continue to solicit and receive food from their parents until about 40 days old, when they become completely independent (Miller and Miller, 1968; Tubaro, 1990). Accordingly, individuals captured during November (spring) would correspond to late September broods, while those captured in mid-January (summer) would correspond to early December broods, ensuring that both broods were raised under markedly different ecological conditions.

Immediately after capture, individuals were placed in a bird-holding bag (Ecotone, Poland), weighed to the nearest 0.1 g on a digital scale, and a ∼50μl blood sample was collected from the brachial vein using heparinized microhematocrit capillary tubes. We then sampled claw tips, which were placed in an Eppendorf tube. All samples were stored frozen in liquid nitrogen prior to preparation for stable isotope analysis. All tissue samples were dried at 80°C, homogenized, and ∼0.5–0.6 mg of each sample was loaded into tin capsules for stable isotope analysis.

### Resource Availability

The rufous-collared sparrow is an omnivorous species that routinely consumes arthropods, seeds, and small herbaceous plants (Lopez-Calleja, 1990, 1995). In order to support the information given by previous research by Lopez-Calleja (1995) at the study site, we examined seasonal variation in resource availability measured as the abundance and diversity of herbaceous plants and terrestrial arthropods. Herbaceous plants were sampled in five permanent 1m^2^ quadrats distributed across the study area which covered approximately 1000 m² within a ravine, ensuring each quadrat was positioned at a minimum distance of 35 meters apart. The greatest distance between two quadrats was 140 meters. On each quadrat, we identified the percent cover of living herbaceous species during both seasons: mid-September and mid-October for spring and mid-December and mid-January for summer. Also, we sampled terrestrial arthropods with Barber pitfall traps filled with water and biodegradable detergent (Southwick, 1978). They were placed every 1 m along five linear transects of 10 m distributed across the study area, at close proximity from the vegetation quadrats, and collected after 24 h (Jaksic and Lazo, 1999). We counted and identified the collected arthropods and classified them into morphotypes corresponding to the lowest taxonomic resolution possible. We then estimated the mean total abundance per sample unit for herbaceous plants and terrestrial arthropods for each season and its confidence interval by bootstrap procedure with 10,000 iterations.

Resource diversity was estimated by quantifying the Hill numbers for the plant and arthropod assemblages (Chao et al., 2014). Seasonal species richness and Shannon diversity index were calculated by using sample-size-based rarefaction and extrapolation estimation of the Hill numbers of order q equal to 0 and 1, respectively. The estimates were based on species abundance, measured as the percent cover of living plant species and number of individuals within each arthropod morphotype (Chao et al., 2014; Hseih et al., 2019). All estimations were performed with the package iNEXT (Hseih et al., 2019).

Moreover, we collected leaf and seed samples from 12 plant species during the spring and seed samples from 6 species during the summer, as live plant cover was absent in our study quadrats. The plant and seed species were selected based on their relative abundance and their inclusion in the rufous-collared sparrow’s diet as described by Lopez-Calleja (1990,1995). This collection allowed us to characterize the δ^15^N composition of the trophic web baseline required to estimate relative trophic level (see below).

### Stable Isotope Analysis

We measured δ^15^N values in blood plasma, red blood cells, and claw tips of juvenile and adult individuals to quantify niche characteristics in seasons characterized by high (spring) and low (summer) resource availability according to previous studies (Lopez-Calleja, 1995). Although many tissues can be compared to construct a time series of isotopic variation, blood plasma, red blood cells, and claws can be easily collected in the field without sacrificing animals (Podlesak et al., 2005; Carleton et al., 2008). δ^15^N values of claw tips integrate dietary inputs over ∼6 weeks prior to sample collection, while blood plasma and red blood cells integrate diet over ∼1-2 and ∼3-4 weeks prior to collection, respectively (Hobson and Clark, 1992; Carleton et al., 2008; Hahn et al., 2014). Young birds initiate the post-juvenile molt either at the time of independence or within about 10 days thereafter (Miller and Miller, 1968). In our study, the captured juveniles showed no evidence of molting, which suggest that probably were not yet fully independent in their feeding. This implies that the variance in diet composition based on analysis of these three tissues, with the most recent one reflecting diet integrated over ∼1-2 weeks prior to capture, represents mostly the diet of juvenile individuals during the parental care period when they had not yet achieved foraging independence.

Nitrogen isotope (δ^15^N) values of sparrow tissues were measured on a Costech 4010 elemental analyzer coupled to a Thermo Finnigan Delta Plus XP isotope ratio mass spectrometer at the University of New Mexico Center for Stable Isotopes (Albuquerque, NM). Stable isotope data are expressed in delta (δ) notation as parts per thousand (‰) according to the equation, δ^15^N = (R_sample_ / R_standard_ – 1) X 1000, where R represents the ratio heavy to light isotopes (^15^N/^14^N) of the sample relative to the international reference standard. Measured δ^15^N values were calibrated relative to air with certified reference materials. Within-run estimates of analytical precision were obtained by measurements of an internal laboratory reference material and yielded a precision (SD) ≤ 0.2‰.

### Total Niche Width and Individual Diet Specialization

To quantify the total niche width and degree of individual diet specialization between age groups and seasons, we used a modified version of Roughgarden index for use with δ^15^N data (Roughgarden, 1972; Maldonado et al., 2017, 2019). Roughgarden used a simple framework to quantify individual diet specialization: the total niche width (TNW) of the (adult or juvenile) age group is the sum of two components: (1) the within-individual component (WIC) which reflects the average of the variability of resources used by individuals, and (2) the between individual component (BIC) that represents the between-individual variation in the average use of resources. The degree of individual diet specialization is reflected in the WIC/TNW ratio where low ratios close to zero indicate a higher prevalence of specialization within an age class, indicating a generalist population composed of individual specialists that each use a small subset of the total niche width.

### Trophic Level

In food web studies involving animals, δ^15^N measurements are commonly used to infer the relative trophic position of consumers due to the predictable and significant enrichment of ^15^N at higher trophic levels, known as trophic enrichment factors (TEFs). This enrichment occurs because of protein catabolism. During catabolism, amino acids with amine groups containing the heavier ^15^N isotope are preferentially retained over those with the lighter ^14^N isotope. Consequently, excreted urinary nitrogen is depleted in ^15^N compared to the animal’s tissues. This leads to a progressive accumulation of ^15^N along the food chain, with each successive trophic level exhibiting higher ^15^N/^14^N ratios than the previous level (Post, 2002; see also Karasov & Martinez del Rio, 2007 and references therein). Additionally, using stable isotopes to estimate trophic levels requires a priori estimates of discrimination factors, which represents the differences in isotopic composition between an animal’s tissue and its diet. These factors are crucial for accurately interpreting δ^15^N data and inferring trophic positions in bird studies, as they vary depending on the tissue being analyzed. We used previously determined discrimination factors for bird tissues in our trophic level calculations (Caut et al.2009). We calculated the relative trophic level (TL) of each individual as follows (Post, 2002): TL= (1 + (δ^15^N_bird_ − δ^15^N_producers_) / Δ^15^N, where the δ^15^N_bird_ represents the mean isotopic signature of tissues, δ^15^N_producers_ correspond to the baseline isotopic signature of the primary producers (plants) and Δ^15^N is the trophic discrimination factor. We used a trophic discrimination of 2.37‰ for collagen (claws), 2.25‰ for blood, and 2.82‰ for plasma (Caut et al., 2009). To calculate the baseline, we estimated the δ^15^N values of plants and seeds for both seasons, but a t-test showed no significant differences between them. Therefore, we used a pooled sample δ^15^N value of 3.47‰ for the baseline at the study site.

### Statistical Analysis

To assess for differences in levels of IS of juveniles and adults against a null model, we employed a non-parametric Monte Carlo procedure with 10,000 iterations to derive p-values. The null model characterizes the total niche width of either juveniles or adults as being comprised of generalist individuals who forage randomly across the entire niche of their respective age classes (Bolnick et al., 2002). This Monte Carlo procedure is implemented in the R package RInSp (Zaccarelli et al., 2013). To evaluate whether significant differences exist in TNW, BIC, WIC, and WIC/TNW between juveniles and adults, we also used a Monte Carlo permutation procedure. To calculate a p-value, the observed mean difference in WIC/TNW between juveniles and adults was compared against the mean differences generated from 10,000 random permutations of juvenile-adult groups. We then assessed potential differences in trophic levels between the two age classes across seasons by linear mixed models (lmm; package lme4, Bates et al., 2015) with age class (juveniles, adults), seasons (spring, summer) and the interaction among them as fixed factors and tissue (nails, blood plasma and red blood cells) as random intercept. Model assumptions (normality, homoscedasticity and absence of residual autocorrelation) were confirmed. Finally, we estimated the isotopic niche overlap between juveniles and adults in both seasons by calculating the overlapped areas (and their bootstrap standard errors with 1000 replicates, SE) relative to each pair of δ^15^N kernel density estimations using the overlapping R package (Pastore, 2018; Pastore and Calcagnì, 2019). To assess whether the estimated area of overlap was significantly different from what would be expected by chance -due solely to sample size-we performed a permutation test with 10,000 random permutations of the δ^15^N values. We then compared the observed overlap area to the 95% confidence interval (CI_0.95_) generated from the permuted data.

## Results

The herbaceous percent plant cover decreased from 86% (CI_0.95_: 82.1 - 90.1%) in spring to 0% during the summer. A total of 27 herbaceous plant species were available during spring, with an estimated Shannon diversity (D^q=1^) of 4.5 (CI_0.95_: 4.3 - 5.3). Conversely, no significant differences were observed in arthropod mean total abundance between seasons; mean= 20.7 (CI_0.95_: 9.7 - 37.7) in spring and mean= 15.2 (CI_0.95_: 9.4 - 22.6) in summer. Also, the Shannon diversity index for arthropods did not differ significantly for spring (D^q=1^ = 11.3; CI_0.95_: 10.9 - 11.9) and summer (D^q=1^ = 12.4; CI: 11.9 - 13.6; See Supplementary Materials). However, the taxonomic richness of arthropods was significantly higher in spring (D^q=0^ = 132, CI_0.95_: 120.9 - 143) than in summer (D^q=0^ = 81, CI_0.95_: 73.6 - 88.4).

Individual diet specialization (IS) and total niche width (TNW) varied significantly between seasons for juveniles but not for adults (Figure 1, Table 1). Juveniles exhibited higher degrees of IS in spring than in summer, while TNW was higher in summer than in spring. In contrast, adults maintained a consistent degree of IS and TNW across seasons. Notably, adults showed a decrease in trophic level in summer relative to spring, while juveniles maintained a consistent trophic level between seasons, which was shown by a significant interaction effect between age and season in our model (β_df=159_ = -0.75, t = - 3.34, p = 0.001; Figure 2, Table S1). Moreover, the amount of niche overlap between juveniles and adults varied from 0.59 ± 0.09 in spring to 0.39 ± 0.09 in summer, being significantly smaller than what would be expected by chance in summer (permuted 95% quantiles: 0.49 - 0.89).

**Figure 1:**
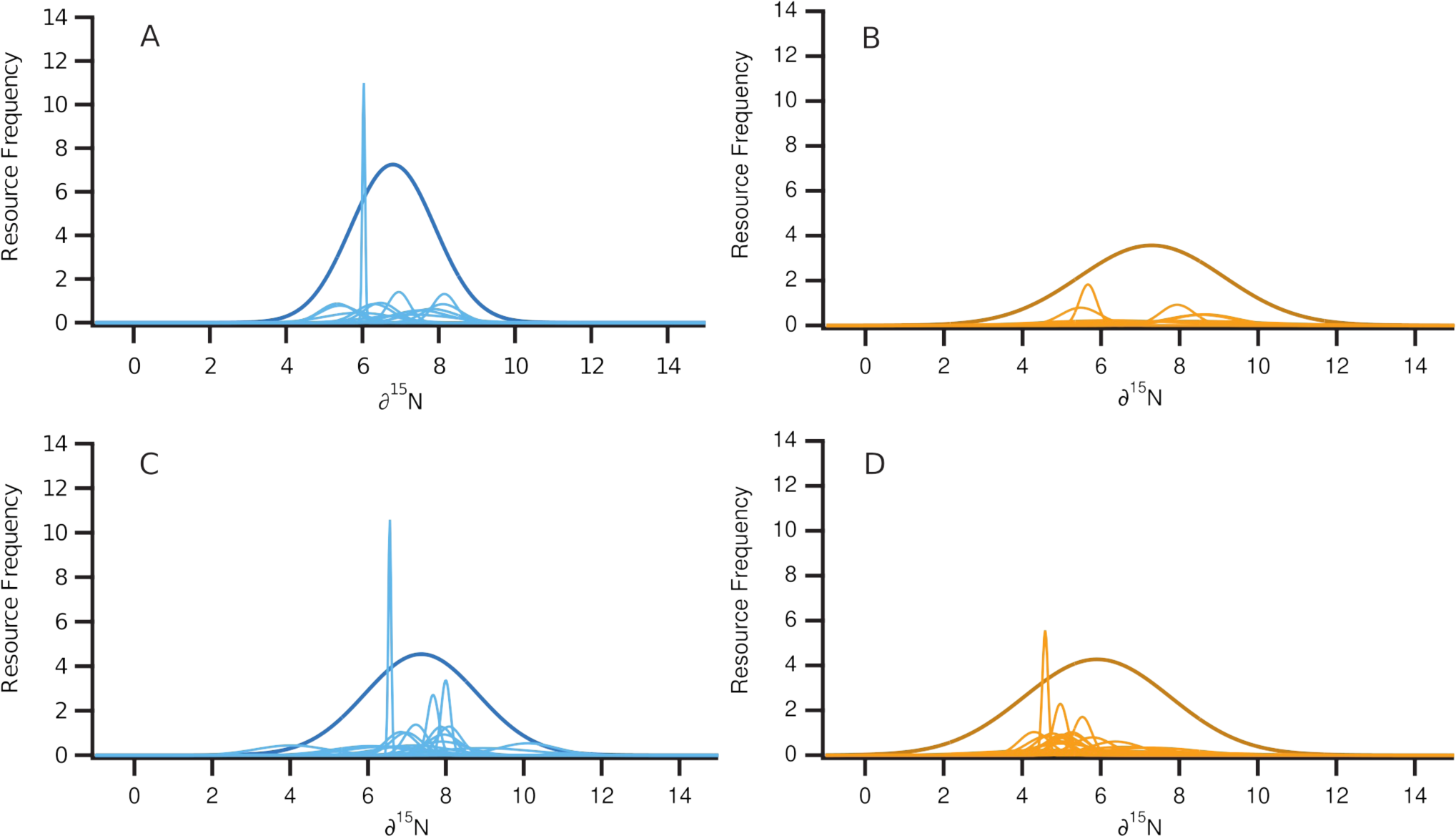
Distributions with thin lines correspond to individual niches for sparrows captured in the spring (blue) and summer (orange) of juveniles (A and B) and adults (C and D) generated from δ^15^N values of blood plasma, red blood cells, and claw tips. Distributions with thick lines represent the corresponding total niche width (TNW), which were scaled equally for the four groups enabling direct comparison.

**Figure 2:**
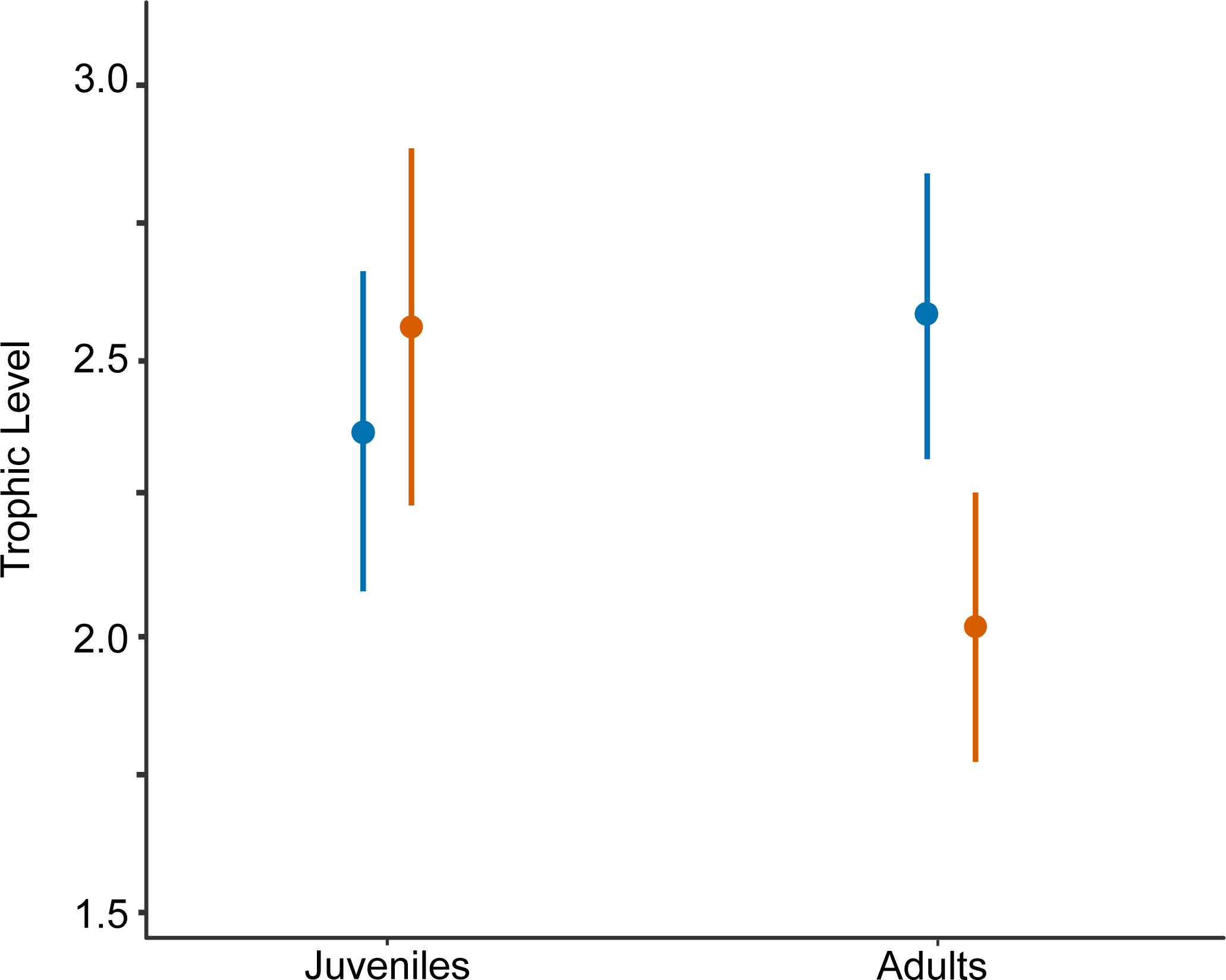
The figure displays the trophic levels of juvenile and adult sparrows during spring (blue) and summer (orange), calculated using δ^15^N isotopic values from diet and bird tissues. The error bars represent the 95% confidence intervals.

**Table 1:** Estimated within-individual component (WIC), between-individual component (BIC), total niche width (TNW), and individual specialization (WIC/TNW) of adults and juveniles individual rufous-collared sparrows captured during summer and spring. Units of WIC, BIC, and TNW are ‰^2^; sample sizes are noted in parentheses.

|  | Spring |  | Summer |  |
| --- | --- | --- | --- | --- |
|  | Juveniles (11) | Adults (16) | Juveniles (8) | Adults (20) |
| **WIC** | 0.252<sup>a</sup> | 0.436<sup>a</sup> | 1.924<sup>b</sup> | 0.566<sup>a</sup> |
| **BIC** | 0.958<sup>a</sup> | 1.665<sup>a</sup> | 1.482<sup>a</sup> | 2.980<sup>a</sup> |
| **TNW** | 1.210<sup>a</sup> | 2.101<sup>a</sup> | 3.405<sup>b</sup> | 3.546<sup>a</sup> |
| **WIC/TNW** | 0.208<sup>a</sup> | 0.207<sup>a</sup> | 0.565<sup>b</sup> | 0.160<sup>a</sup> |

Different superscript letters denote significant differences ( $P < 0.05$ ) among groups.

## Discussion

This study offers valuable insights into the trophic niche structure and individual specialization (IS) of juveniles and adults during periods of varying resource availability in a species that provides parental care both in the nest and during the post-fledging period. Contrary to our expectations, we found that resource availability influences both total niche width (TNW) and the degree of IS in juveniles (Table 1). However, adults showed no change in these niche characteristics over the breeding season despite a shift in their trophic level (Figure 2).

There are two primary hypotheses that relate resource availability to niche subdivision and IS within populations. The Niche Variation Hypothesis (NVH) suggests that an increase in the quantity and diversity of resources allows individuals to specialize in different resources, broadening the overall TNW and increasing IS (Van Valen, 1965; Bolnick et al., 2002, 2003). On the other hand, the Optimal Foraging Theory (OFT) posits that when resource availability increase, animals will concentrate on the most preferred and profitable resource (Stephens and Krebs, 1986; Araújo et al., 2011). This leads to a narrower TNW and reduced IS, as more individuals exploit the same optimal resources (Newsome et al., 2015; Manlick et al., 2021). These seemingly opposing hypotheses have both found support in various ecological studies, indicating that the relationship between resource availability and niche dynamics can be complex and context-dependent. Our findings, supported by previous studies (Lopez-Calleja, 1995), indicate that resource availability is highest in spring due to increased arthropod richness and greater plant cover and diversity. According to the NVH, the greater availability of resources would enable adults to specialize their resource use for them and their offspring. Consequently, both age classes were primarily composed of individual specialists, which collectively exploit a wide range of resources, exhibiting nearly identical degrees of IS, TNW and trophic levels. The high niche overlap found between juveniles and adults further indicates the similarities in niche characteristics between these two groups during this season. Following the same rationale, we would expect that in age classes faced with a reduction in food resources, the TNW and degree of IS should decrease for both adults and juveniles. In contrast to this prediction, the prevalence of IS only decreased in juveniles in summer by means of a slight increase of WIC and a broadening of their TNW (Table 1). Our explanation is that juveniles require a protein-rich diet for development, which in turn compels adults to provide that kind of food to maintain juvenile requirements under conditions of low resource availability potentially at the cost of juvenile IS levels. This explanation aligns with OFT, suggesting that parents feed their offspring with additional less preferred food items when resources are scarce, which results in a wider TNW and more generalist individual niches observed in juveniles during the summer relative to spring, while maintaining the trophic level.

The TNW and IS of adults were similar across seasons, but this age class occupied a lower trophic level in summer relative to spring. This resulted in a significantly lower amount of dietary overlap between juveniles and adults during the summer. This suggests that adults may be inherently more flexible in resource consumption because they are not constrained by the same nutritional requirements as developing nestlings. These conditions enable adults to maintain their levels of IS during periods of relative resource scarcity while prioritizing the delivery of higher trophic level prey items to their nestlings during critical periods of their growth and development.

In conclusion, the results of this study highlight a dynamic interplay between resource availability and IS, with significant implications for understanding species resilience to environmental variation (Bolnick et al., 2003; Araújo et al., 2011). Notably, our study reveals that adult sparrows modify nest provisioning strategies in response to seasonal fluctuations in resource availability. For juveniles, the shift towards generalism in times of resource scarcity may be imperative to survival, ensuring adequate nutrient intake for growth and development. In contrast, the adult’s ability to maintain niche width but alter trophic position underlines a strategic flexibility that likely minimizes intraspecific competition and optimizes resource utilization across the population. Ultimately, this study underscores the importance of considering IS to fully understand the ecological and evolutionary mechanisms governing niche partitioning among and within populations. In Mediterranean ecosystems like central Chile, climate change projections predict an increase in temperatures and a decrease in annual precipitation (Bozkurt et al., 2017, 2018), therefore the relatively dry resource-limited conditions associated with summer may extend across a larger portion of the reproductive season. Studies like ours that focus on characterizing dietary flexibility and understanding how resource use may shift under these conditions are essential for understanding species response(s) to ongoing environmental change.

## References

Araújo, M. S., Bolnick, D. I., and Layman, C. A. (2011). The ecological causes of individual specialisation. Ecology Letters 14, 948–958. doi: 10.1111/j.1461-0248.2011.01662.x

Arco, L., Martín-Vivaldi, M., Peralta-Sánchez, J. M., Juárez García-Pelayo, N., and Soler, M. (2022). Provisioning challenge: self-consumption versus nestling provisioning, an experimental study. Animal Behaviour 190, 153–165. doi: 10.1016/j.anbehav.2022.06.008

Bates, D., Maechler, M., Bolker, B., Walker, S., Christensen, R. H. B., Singmann, H.,… & Bolker, M. B. (2015). Package ‘lme4’. convergence, 12(1), 2.

Beaulieu, M., and Sockman, K. W. (2014). Comparison of optimal foraging versus life-history decisions during nestling care in Lincoln’s Sparrows *Melospiza lincolnii* through stable isotope analysis. Ibis 156, 424–432. doi: 10.1111/ibi.12146

Bolnick, D. I., Svanbäck, R., Fordyce, J. A., Yang, L. H., Davis, J. M., Hulsey, C. D., et al. (2003). The ecology of individuals: Incidence and implications of individual specialization. American Naturalist 161, 1–28. doi: 10.1086/343878

Bolnick, D. I., Yang, L. H., Fordyce, J. A., Davis, J. M., and Svanbäck, R. (2002). Measuring individual-level resource specialization. Ecology 83, 2936–2941.

Bozkurt, D., Rojas, M., Boisier, J. P., and Valdivieso, J. (2017). Climate change impacts on hydroclimatic regimes and extremes over Andean basins in central Chile. Hydrology and Earth System Sciences Discussions, 1–29. doi: 10.5194/hess-2016-690

Bozkurt, D., Rojas, M., Boisier, J. P., and Valdivieso, J. (2018). Projected hydroclimate changes over Andean basins in central Chile from downscaled CMIP5 models under the low and high emission scenarios. Climatic Change 150, 131–147. doi: 10.1007/s10584-018-2246-7.

CR2. Center for Climate and Resilience Research, Chile.

Carleton, S. A., Kelly, L., Anderson-Sprecher, R., and del Rio, C. M. (2008). Should we use one-, or multi-compartment models to describe 13C incorporation into animal tissues? Rapid Communications in Mass Spectrometry: An International Journal Devoted to the Rapid Dissemination of Up-to-the-Minute Research in Mass Spectrometry 22, 3008–3014.

Chao, A., Gotelli, N. J., Hsieh, T. C., Sander, E. L., Ma, K. H., Colwell, R. K., et al. (2014). Rarefaction and extrapolation with Hill numbers: a framework for sampling and estimation in species diversity studies. Ecological Monographs 84, 45–67. doi: 10.1890/13-0133.1

Di Castri, F., and Hajek, E. R. (1976). Bioclimatología de chile.

Durell, S. leV Dit. (2003). The implications for conservation of age-and sex-related feeding specializations in shorebirds. Wader Study Group Bulletin, 35–39.

Finke, D. L., and Snyder, W. E. (2008). Niche Partitioning Increases Resource Exploitation by Diverse Communities. Science 321, 1488–1490. doi: 10.1126/science.1160854

Frédérich, B., Lehanse, O., Vandewalle, P., and Lepoint, G. (2010). Trophic niche width, shift, and specialization of *Dascyllus aruanus* in Toliara Lagoon, Madagascar. Copeia, 218–226. doi: 10.1643/CE-09-031

Hahn, S., Dimitrov, D., Rehse, S., Yohannes, E., and Jenni, L. (2014). Avian claw morphometry and growth determine the temporal pattern of archived stable isotopes. Journal of Avian Biology 45, 202–207. doi: 10.1111/j.1600-048X.2013.00324.x

Hall, L. A., De La Cruz, S. E. W., Woo, I., Kuwae, T., and Takekawa, J. Y. (2021). Age- and sex-related dietary specialization facilitate seasonal resource partitioning in a migratory shorebird. Ecology and Evolution 11, 1866–1876. doi: 10.1002/ece3.7175

Hammerschlag-Peyer, C. M., Yeager, L. A., Araújo, M. S., and Layman, C. A. (2011). A hypothesis-testing framework for studies investigating ontogenetic niche shifts using stable isotope ratios. PLoS ONE 6. doi: 10.1371/journal.pone.0027104

Hobson, K. A. (1993). Trophic relationships among high Arctic seabirds: insights from tissue-dependent stable-isotope models. Marine Ecology Progress Series 95, 7–18. doi: 10.3354/meps095007

Hobson, K. A., and Clark, R. G. (1992). Assessing avian diets using stable isotopes I: Turnover of 13C in tissues. Condor 94, 181–188. doi: 10.2307/1368807

Hseih, T., Ma, K., and Chao, A. (2019). iNEXT: iNterpolation and EXTrapolation for species diversity. R package 2, 19.

Jaksic, F. M., and Lazo, I. (1999). Response of a bird assemblage in semiarid Chile to the 1997-1998 El Niño. The Wilson Bulletin, 527–535.

Jia, G., Shevliakova, E., Artaxo, P., Noblet-Ducoudré, N. D., Houghton, R., House, J., et al. (2019). “Land–climate interactions,” in Climate Change and Land: an IPCC special report on climate change, desertification, land degradation, sustainable land management, food security, and greenhouse gas fluxes in terrestrial ecosystems.

Kyriazakis, I., and Oldham, J. D. (1993). Diet selection in sheep: the ability of growing lambs to select a diet that meets their crude protein (nitrogen × 6.25) requirements. British Journal of Nutrition 69, 617–629. doi: 10.1079/bjn19930064

López-Calleja, M. V. (1990). Variacion estacional en el uso de los recursos alimenticios por algunos componentes de una taxocenosis de aves paseriformes en Quebrada de la Plata, Chile central.

Lopez-Calleja, M. V. (1995). Dieta de *Zonotrichia capensis* (Emberizidae) Y *Diuca diuca* (Fringillidae): efecto de la variación estacional de los recursos tróficos y la riqueza de aves granívoras en Chile central. Revista Chilena de Historia Natural 68, 321–331.

Maldonado, K., Bozinovic, F., Newsome, S. D., and Sabat, P. (2017). Testing the niche variation hypothesis in a community of passerine birds. Ecology 98, 903–908. doi: 10.1002/ecy.1769

Maldonado, K., Newsome, S. D., Razeto-Barry, P., Ríos, J. M., Piriz, G., and Sabat, P. (2019). Individual diet specialisation in sparrows is driven by phenotypic plasticity in traits related to trade-offs in animal performance. Ecology Letters 22, 128–137. doi: 10.1111/ele.13174

Manlick, P. J., Maldonado, K., and Newsome, S. D. (2021). Competition shapes individual foraging and survival in a desert rodent ensemble. Journal of Animal Ecology 90, 2806– 2818. doi: 10.1111/1365-2656.13583

Miller, A. H., and Miller, V. D. (1968). the Behavioral Ecology and Breeding Biology of the Andean Sparrow, Zonotrichia Capensis. Source: Caldasia 10, 83–154.

Newsome, S. D., Tinker, M. T., Gill, V. A., Hoyt, Z. N., Doroff, A., Nichol, L., et al. (2015). The interaction of intraspecific competition and habitat on individual diet specialization: a near range-wide examination of sea otters. Oecologia 178, 45–59. doi: 10.1007/s00442-015-3223-8

Pastore, M. (2018). Overlapping: a R package for Estimating Overlapping in Empirical Distributions. Journal of Open Source Software 3, 1023. doi: 10.21105/joss.01023

Pastore, M., and Calcagnì, A. (2019). Measuring Distribution Similarities Between Samples: A Distribution-Free Overlapping Index. Frontiers in Psychology 10.

Podlesak, D. W., McWilliams, S. R., and Hatch, K. A. (2005). Stable isotopes in breath, blood, feces and feathers can indicate intra-individual changes in the diet of migratory songbirds. Oecologia 142, 501–510.

Ramos, R., Reyes-González, J. M., Morera-Pujol, V., Zajková, Z., Militão, T., and González-Solís, J. (2020). Disentangling environmental from individual factors in isotopic ecology: A 17-year longitudinal study in a long-lived seabird exploiting the Canary Current. Ecological Indicators 111, 105963. doi: 10.1016/j.ecolind.2019.105963

Riverón, S., Raoult, V., Baylis, A. M. M., Jones, K. A., Slip, D. J., and Harcourt, R. G. (2021). Pelagic and benthic ecosystems drive differences in population and individual specializations in marine predators. Oecologia 196, 891–904. doi: 10.1007/s00442-021-04974-z

Roughgarden, J. (1972). Evolution of niche width. The American Naturalist 106, 683–718.

Southwick, L. H. (1978). Ecological methods: with particular reference to the study of insect populations. Chapman and Hall.

Stearns, S. C. (1998). The evolution of life histories. Oxford university press.

Stephens, D. W., and Krebs, J. R. (1986). Foraging theory. Princeton university press.

Stephens, R. B., Hobbie, E. A., Lee, T. D., and Rowe, R. J. (2019). Pulsed resource availability changes dietary niche breadth and partitioning between generalist rodent consumers. Ecology and Evolution 9, 10681–10693. doi: 10.1002/ece3.5587

Sword, G. A., and Dopman, E. B. (1999). Developmental specialization and geographic structure of host plant use in a polyphagous grasshopper, *Schistocerca emarginata* (=lineata) (Orthoptera: Acrididae). Oecologia 120, 437–445. doi: 10.1007/s004420050876

Tubaro, P. L. (1990). Aspectos causales y funcionales de los patrones de variación del canto del chingolo (*Zonotrichia capensis*). Universidad de Buenos Aires. Facultad de Ciencias Exactas y Naturales.

Van Valen, L. (1965). Morphological variation and width of ecological niche. The American Naturalist 99, 377–390.

Vander Zanden, H. B., Bjorndal, K. A., and Bolten, A. B. (2013). Temporal consistency and individual specialization in resource use by green turtles in successive life stages. Oecologia 173, 767–777. doi: 10.1007/s00442-013-2655-2

Zaccarelli, N., Bolnick, D. I., and Mancinelli, G. (2013). RI n S p: an r package for the analysis of individual specialization in resource use. Methods in Ecology and Evolution 4, 1018–1023.

