## Supplementary Material for "The Interplay of Resource Availability and Parent Foraging Strategies on Juvenile Sparrow Individual Specialization"

**Diversity of Arthropods**

Shannon diversity for arthropods was estimated as the asymptotic Hill number of order q = 1. However, species richness estimation did not reach the asymptote for an extrapolated sample size below the triple minimum reference sample size (N = 1503) (see Figure 1). Following the recommendations of Chao et al. (2014) and to enable a precise comparison of species richness across seasons, Hill numbers of order q = 0 were estimated for sample sizes ranging from the double minimum reference sample size to the maximum reference sample size (N = 3458), ensuring consistent sample coverage– a metric used to measure the completeness of a sample, that represents the proportion of individuals that belong to the observed species in a sample (Figure 2) –.

The results reveal a significantly higher richness of arthropods in spring compared to summer. For richness estimation, the reference sample size (N = 1503 for summer and N = 3458 for spring) represent a sample coverage of 98.0% for both seasons (Figure 2). At these sample sizes, richness is estimated in 81.00 (CI: 73.60 – 88.40) for summer and 132.00 (CI: 120.94 – 143.06) for spring, with non-overlapping confidence intervals (Figure 1). Figure 1 also illustrates that the difference in species richness increases with sample size, despite similar sample coverage (Figure 2). However, Shannon diversity is estimated as 12.43 (CI: 11.94 – 13.64) for summer and 11.255 (CI: 10.89 – 11.97) for spring, with overlapping confidence intervals, thereby not ensuring a significant difference between both seasons.

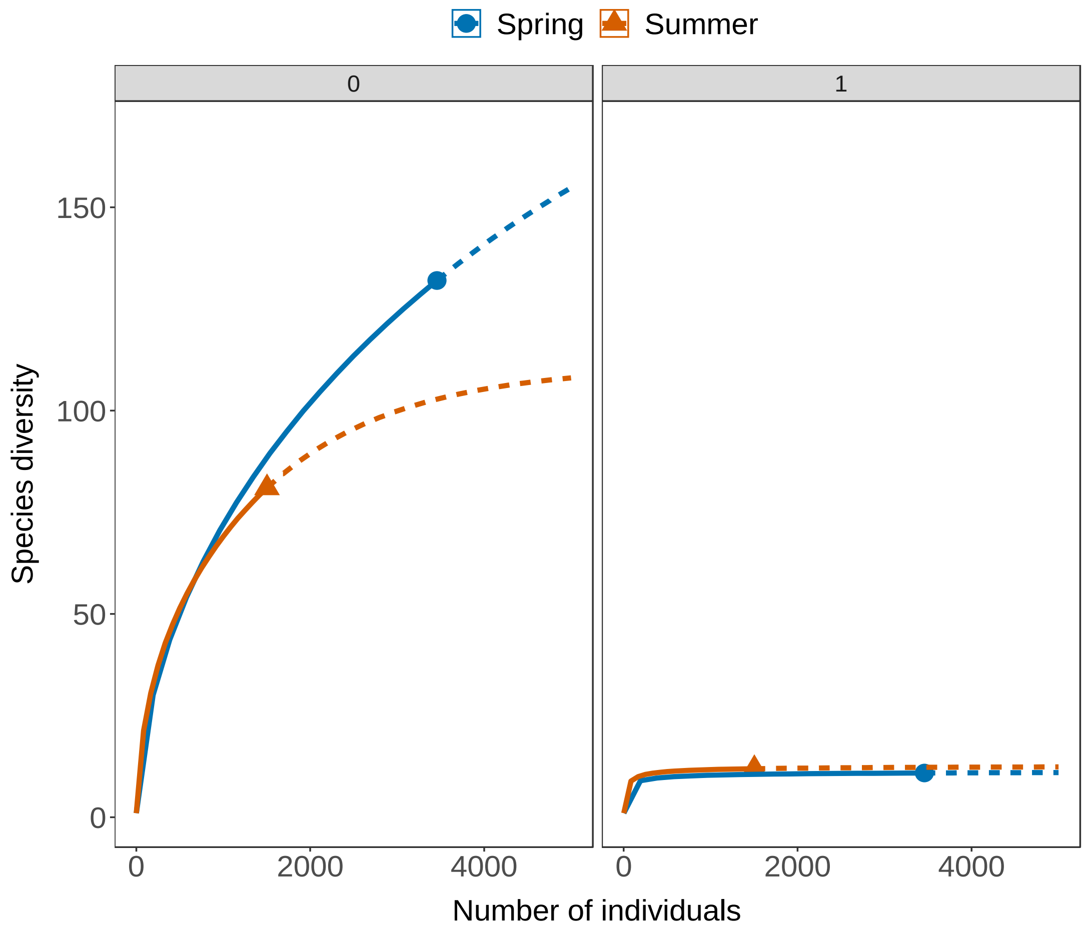

**Figure 1:** Sample-size-based rarefaction (solid lines) and extrapolation (dashed lines) of arthropods diversity based on the Hill numbers (q = 0, 1) for each sampling season. 95% confidence intervals (shaded regions) were obtained by a bootstrap method based on 200 replications. Reference samples are denoted by solid dots.

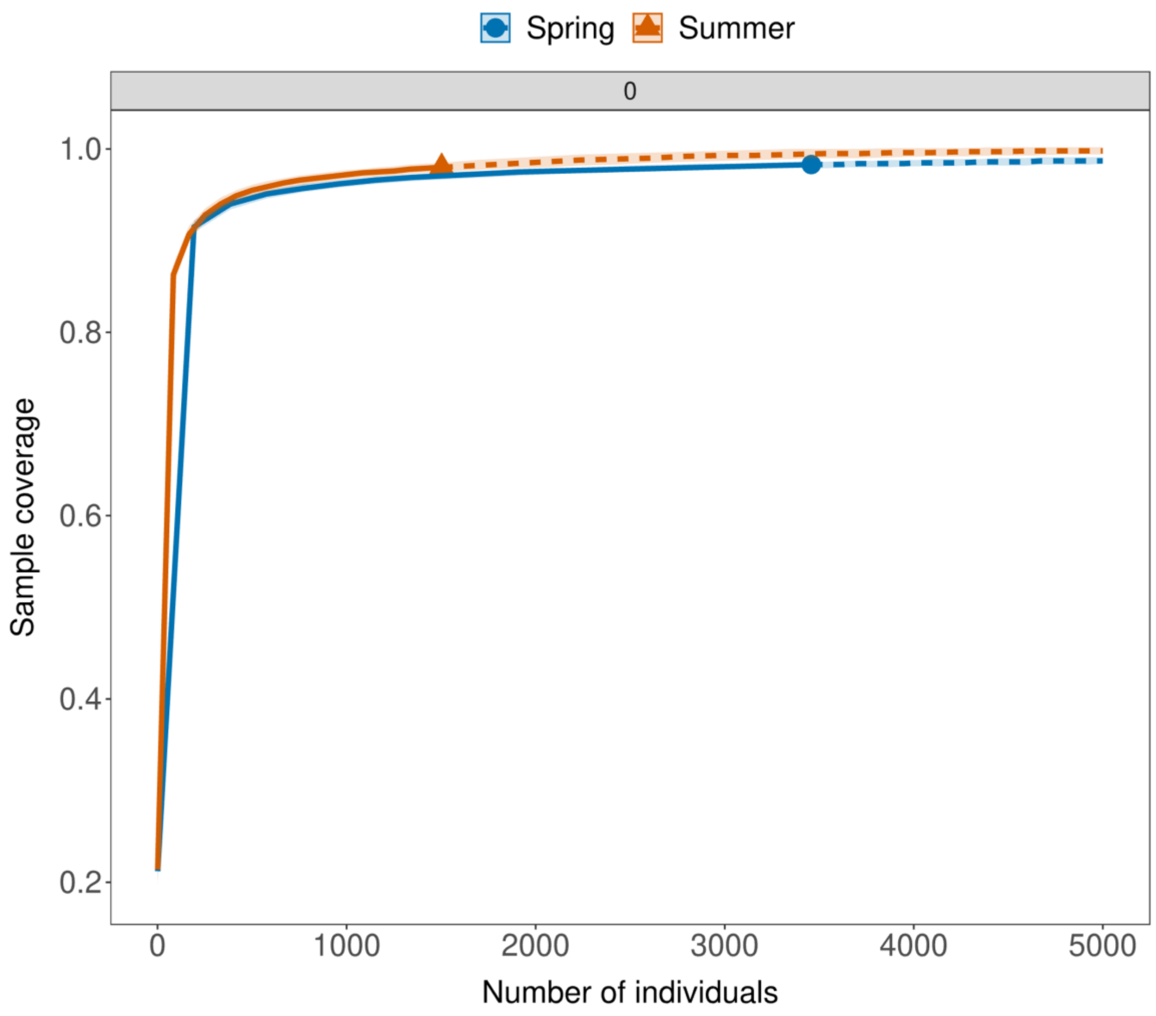


**Figure 2:** Sample coverage for rarefied samples (solid lines) and extrapolated samples (dashed line) with 95% confidence intervals for arthropods sampled at each season. Reference samples are denoted by solid dots.

**Table S1:** Table showing the fixed effects estimates from the mixed model, including standard errors, t-values, degrees of freedom and p-values. Significant effects ($p < 0.05$) are in bold.

| | Estimate (β) | Std. Error | df | t-value | Pr(>\|t\|) |
|---|---|---|---|---|---|
| (Intercept) | 2.373 | 0.147 | 8.5849 | 16.146 | 1.01E-07 |
| Age | 0.208 | 0.154 | 159 | 1.353 | 0.178 |
| Season | 0.189 | 0.183 | 159 | 1.036 | 0.302 |
| Age:Season | -0.753 | 0.225 | 159 | -3.341 | **0.001** |
